# A Non-invasive Detection of Parkinson’s Disease using PitArray: An Integrative Meta-Analysis and Machine Learning Approach

**DOI:** 10.1101/2023.10.15.562440

**Authors:** Arittra Bhattacharjee, Tabassum Binte Jamal, Ishtiaque Ahammad, Anika Bushra Lamisa, Md. Shamsul Arefin, Zeshan Mahmud Chowdhury, Mohammad Uzzal Hossain, Keshob Chandra Das, Chaman Ara Keya, Md Salimullah

**Affiliations:** Bioinformatics Division, National Institute of Biotechnology, Ganakbari, Ashulia, Savar, Dhaka, 1349, Bangladesh; Molecular Biotechnology Division, National Institute of Biotechnology, Ganakbari, Ashulia, Savar, Dhaka, 1349, Bangladesh; Department of Biochemistry and Microbiology, North South University, Bashundhara, Dhaka, 1229, Bangladesh

**Keywords:** Parkinson’s disease, microarray, meta-analysis, differential gene expression (DEGs), machine learning, deep learning

## Abstract

Parkinson’s disease (PD) is a progressive neurodegenerative disorder affecting the central nervous system, often diagnosed in its advanced stages due to the absence of sensitive biomarkers. With this objective in mind, our study conducted a comprehensive analysis of differentially expressed genes (DEGs) sourced from blood-based microarray datasets to uncover potential biomarkers and developed a machine learning based classifier to conduct two step validations. By analyzing gene expression of three projects, we identified 678 DEGs, consisting of 337 genes showing upregulation and 341 genes presenting downregulation. Additionally, insights from functional enrichment and the protein-protein network analysis indicate that *HLA-F*, *IRF-1*, and *RPS28* have the potential to serve as biomarkers for diagnosing PD. Simultaneously, we employed feature selection techniques such as Least Absolute Shrinkage and Selection Operator with Cross Validation (LassoCV) followed by Recursive Feature Elimination with Cross Validation (REFCV) to filter our initial dataset of 13,249 genes down to 43 genes, which were subsequently used to train the machine learning-based classifier models. These 43 genes formed the basis for training and testing various machine learning models, including logistic regression, random forest, naive Bayes, k-nearest neighbors, support vector machine, and deep learning based artificial neural networks. Our models demonstrated robust performance, with Support Vector Machine outperforming others by 0.65 accuracy (95%CI: 0.58-0.66), 0.70 AUC-ROC (95%CI: 0.70-0.71) and 0.35 MCC (95%CI: 0.34-0.39). The model was implemented to develop the PitArray tool for non-invasive detection of PD from blood. PitArray is available at: https://github.com/Arittra95/PitArray.

**Key Points:**

- *HLA-F, IRF-1,* and *RPS28* were identified as potential biomarkers for Parkinson’s disease diagnosis.
- Several sophisticated feature selection methods recognized 43 genes which were then used to build a machine learning model.
- A Support Vector Machine based tool named PitArray was developed which could distinguish Parkinson’s disease patients from healthy people based on blood transcriptome data.

## Introduction

Parkinson’s Disease (PD) is the second most common neurodegenerative disorder associated with progressive motor symptoms, cognitive impairment, mental health issues, sleeping problems, pain and sensory disturbances. PD typically affects 2% to 3% of those over 65 years old [1]. However, PD-related physical impairments and fatalities have been rising alarmingly in recent years. For instance, the number of deaths and disabilities associated with PD increased by 81% and >100%, respectively, from 2000 to 2019 [2]. About 500,000 people in the USA have been diagnosed with PD, and some experts estimated that another 500,000 go untreated or received the wrong diagnosis, at a yearly cost of $20 billion USD [3].

PD usually develops in the elderly for various environmental reasons (e.g., continuous exposure to pesticides, herbicides, and heavy metals), lifestyle related influences (caffeine and cigarette smoking) and genetic factors (mutations in PARK genes) [4]. Neuropathological studies revealed that PD causes the neuronal loss of substantia nigra (part of the brain that helps to control body movements) and widespread intracellular α-synuclein protein accumulation [5]. In the early stages of the disease, the ventrolateral substantia nigra starts to lose dopaminergic neurons. Through the progression of disease, the dramatic loss of this type of neurons consequently develop motor symptoms [6], [7]. However, before developing motor symptoms, PD causes several non-motor symptoms such as depression, anxiety and constipation [1]. Individuals with untreated PD face neuropsychiatric disorders, bradykinesia, rigidity and tremor at the early stage of motor symptoms. Eventually PD predisposes them to various social, mental and orthopedic issues [8], [9].

The diagnosis of PD is based on symptoms such as rapid eye movement, sleep behavior disturbance, hyposmia, constipation, and an assessment of typical movement difficulty, psychological or cognitive impairments [10]. However, multiple system atrophy, progressive supranuclear palsy, chorea, dystonia and ataxia also share similar features with PD. Hence, these types of diagnosis are challenging, specially in Low- and Middle-Income Countries (LMICs), due to the lack of expert clinicians or trained personnel [11]. Though Magnetic Resonance Imaging (MRI) and Dopamine Transporter (DaT) scans can be useful in identifying PD, these procedures are expensive and routine use of these tests is discouraged by many practitioners [4]. Molecular investigations demonstrated mutations in several genes that are highly associated with PD. For example, mutations in *PRKN*, *UCH-L1*, *PINK1*, *DJ-1*, and *NR4A2* genes have been observed in PD patients [12], [13]. Hence, these mutations are potential prognostic factors for PD. To diagnose PD, blood gene signatures can leverage the identification of this disease by overcoming the limitations regarding regular clinical examinations [12]. Previously some meta-analysis studies revealed differentially expressed genes in the PD patients that can be potential biomarkers [14]–[16]. However, these studies include tissue based gene expression data beside blood gene expression data. Tissue specific data can provide strong results but it is not suitable for non-invasive diagnosis [12]. Several meta-analyses generated prospective machine learning based classifiers using datasets from different microarray platforms [17], [18]. Although results from different microarray platforms are correlated, data from a single platform could produce more reliable results.

In this study, we processed and analyzed only blood based gene expression datasets of PD patients and healthy individuals. All of these data were generated via the most widely used Affymetrix platform. At first, we tried to identify prospective biomarkers by recognizing differentially expressed genes. Afterwards, we made machine learning based classification models that can classify PD from blood gene signatures. Our two step methods hold the potential to provide strong diagnostic support for PD.

## Methods

Our overarching goal was to identify candidate biomarkers of PD (the focus of this study) using the following workflow (Figure 1).

**Figure 1:**
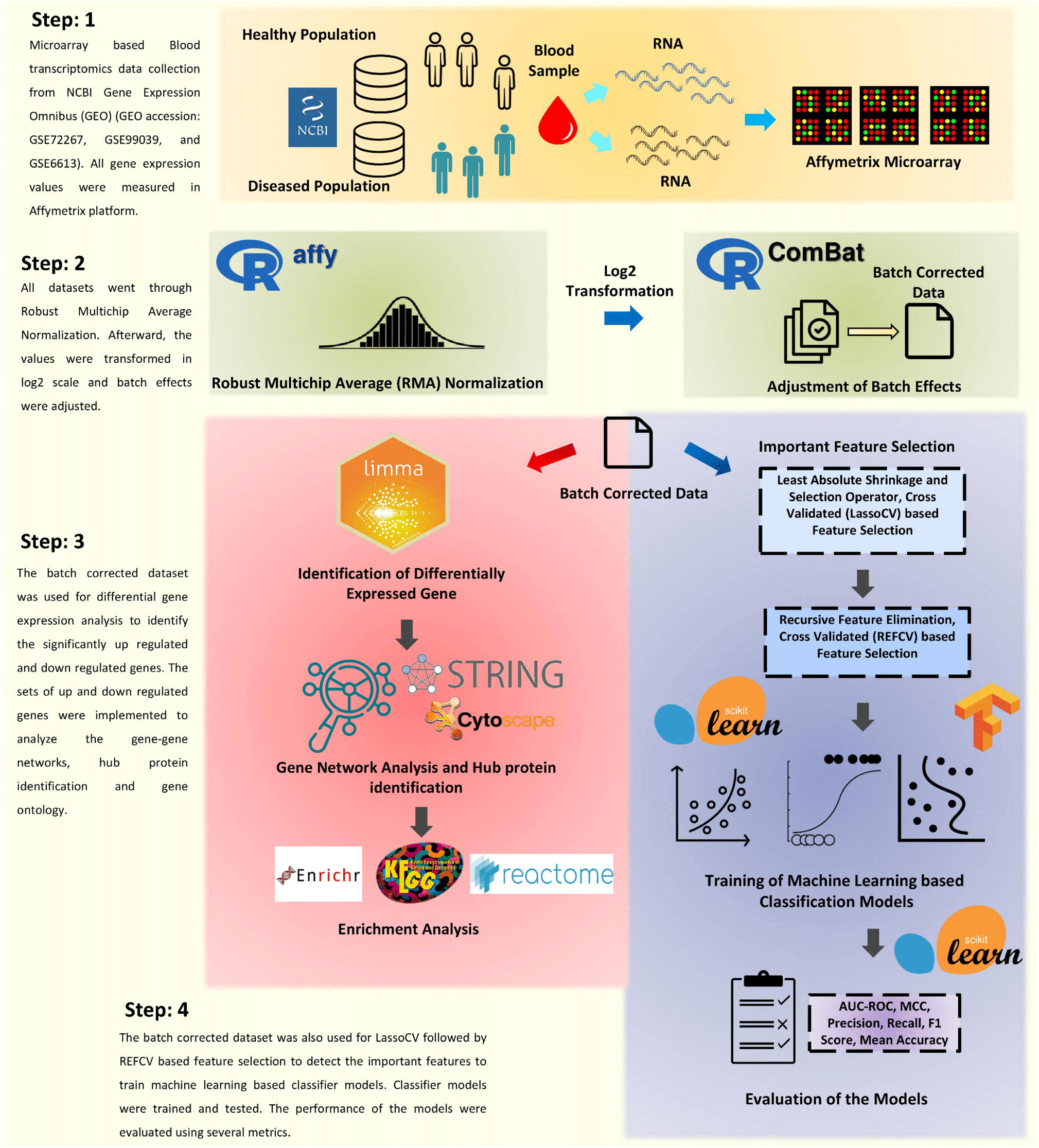
Overall representation of the study. The study was divided into four major steps. In step 1, data collection was conducted. In step 2, the data went through normalization and batch correction. In step 3 (red box), differentially expressed genes were identified and subsequently their network analysis was done with enrichment analysis. Finally, in step 4 (blue box), the data was used to train machine learning models.

### Data collection

The term “Parkinson’s disease” was employed as search keywords in the NCBI-GEO database to locate genome-wide expression studies. The primary focus was on original experimental studies that aimed to identify variations in genes between individuals with PD and healthy individuals. The following aspects were taken into consideration for the inclusion criteria: (1) studies referring to “array-based expression profiling”; (2) availability of raw cell intensity file (CEL) data; and (3) studies involving blood samples. Among the datasets reviewed, three gene expression datasets met these stringent inclusion criteria: GSE72267, GSE99039, and GSE6613. Comprehensive information about these microarray datasets, including details about the platforms used and the number of samples, is provided in Figure 2. It’s important to mention that / notably, some original datasets contained more samples than those utilized in this study. However, certain samples were excluded from analysis due to specific reasons. In the case of dataset GSE99039, 48 patient samples with neurodegenerative disorders were omitted to avoid potential distortions in gene expression patterns. Similarly, 33 samples from the original GSE6613 dataset were also excluded for the same reason.

**Figure 2:**
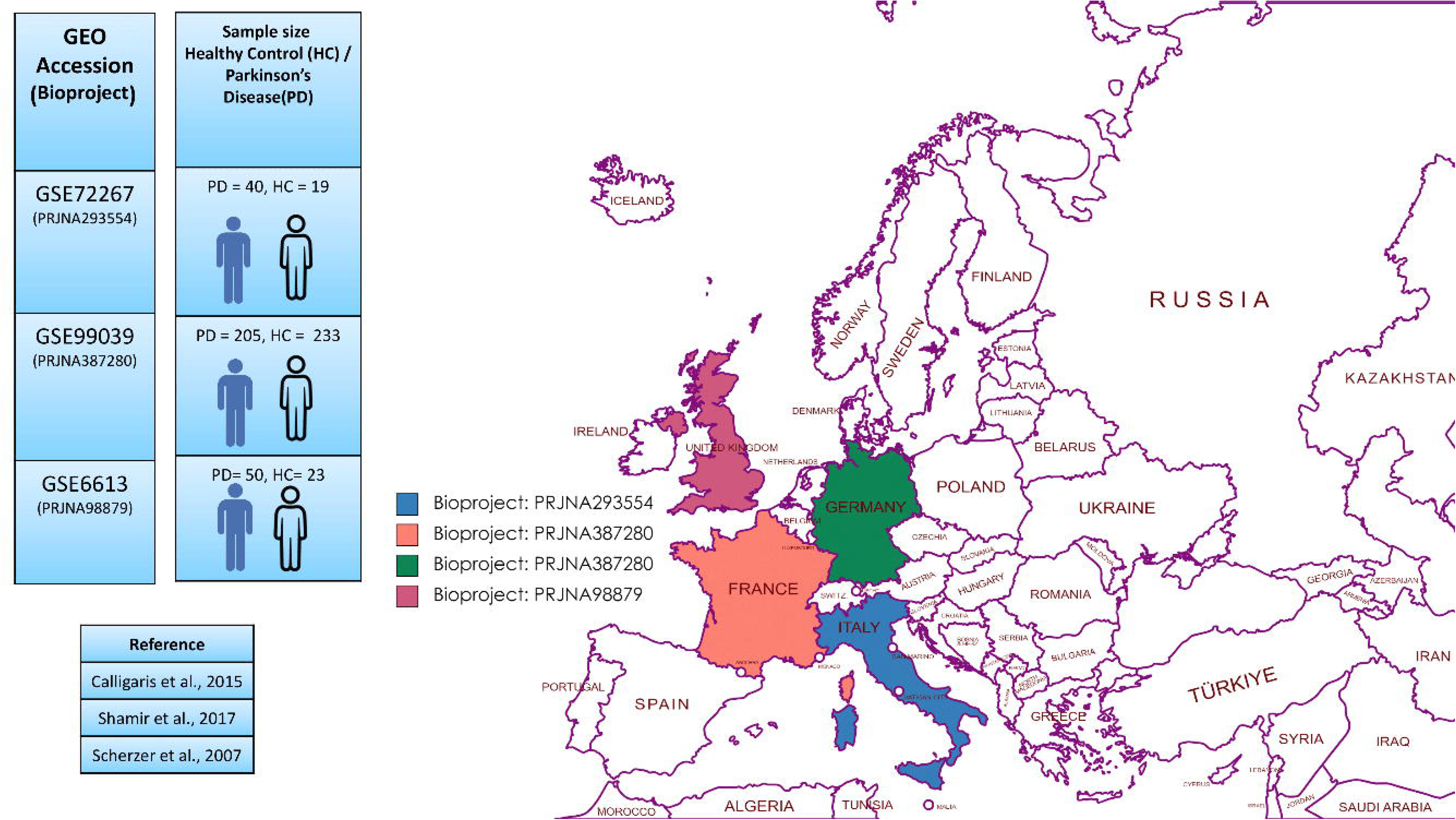
Blood based microarray data was collected from three different studies. The NCBI bioproject numbers of the selected studies have been provided. The studies were conducted in the United Kingdom, Germany, France and Italy.

### Data Preprocessing

Before merging the microarray datasets, the raw data (.cel files) must go through pre-processing. The downloaded data in CEL files were processed using the Affy package (version 1.78.2) in the R language. For individual dataset analysis, each dataset underwent background correction and normalization using the Robust Multichip Averaging (RMA) method [19]. Subsequently, annotations were added to the probes. Probes lacking gene symbol annotations were removed, and if multiple probes were associated with a single gene, the average values were considered as the representative expression value. Afterwards, log transformation was applied to the entire dataset to stabilize the variance of the gene expression data.

### Batch Effect Removal and DEG Screening

Small differences in non-biological factors can affect gene expression results, leading to “batch effects.” These effects stem from variations in time and location during experiments, making samples from different batches not directly comparable. Batch effects are common due to factors like different reagents, timing, and environment, affecting the accuracy of gene expression analyses [20], [21]. To address this issue, “Combating Batch Effects When Combining Batches of Gene Expression Microarray Data” (ComBat) was implemented to adjust the merged gene expression data for batch effects using the ComBat package (0.0.4) in R [21]. Henceforth, to calculate the genes that exhibited differential expression (DEGs) between samples from individuals with PD and those from healthy controls (HC), the limma package (version 3.56.2) was utilized. Subsequently, a comprehensive analysis was conducted, building upon a meticulous selection process that considered adjusted p-values (<0.05), median expression, and average expression thresholds and ultimately leading to the identification of a distinct cohort of differentially expressed genes.

### Functional and Pathway Enrichment Analysis

After combining the initial gene list, the assessment of gene enrichment was performed to reveal functionally associated genes involved in pathways and gene regulation through gene ontology (GO) and Kyoto Encyclopedia of Genes and Genomes (KEGG) pathway enrichment analyses. GO analysis helps to understand the biological significance of gene expression changes within extensive genomic or transcriptomic data. [22]. The significance of changes in gene expression is understood by identifying whether specific GO terms linked to biological processes, molecular functions, or cellular components are disproportionately present in the gene set [23]. Similarly, the KEGG links genomic information with higher-level functional insights by organizing gene annotations and mapping genes to cellular processes [24]. In this study, differentially expressed genes (DEGs) concerning GO terms and KEGG pathways were evaluated using Enrichr [25]. Enrichr is a user-friendly online tool that combines results from various libraries and helps analyze gene lists by providing easy-to-understand visual summaries [26]. Both GO terms and significant pathways were chosen utilizing a Fisher exact test p-value below 0.05 and a high combined score.

### Protein-protein interaction (PPI) network construction for common DEGs

The protein-protein interaction pattern and network (PPI) were obtained using the STRING database [27]. STRING compiles known and predicted protein interactions from various sources, and its latest version (12.0) covers 12535 organisms with 59.3 million proteins, supporting genome-level data uploads. To identify the essential proteins encoded by the DEGs linked to PD, the DEGs were analyzed using STRING, employing a PPI score of 0.4. The generated interaction pattern was downloaded and subjected to further analysis using Cytoscape software (3.10.0). The importance of nodes in the PPI network, like node degree, was studied using the cytoHubba plugin (version 0.1) [28]. Nodes with elevated degrees were identified as hubs. The degree indicates the significance of protein nodes within the network – the higher the degree, the greater the importance of the nodes.

### Feature selection for machine learning models

A two layer hybrid feature selection approach was taken to train the machine learning models from the microarray dataset. To do so, the output dataset of ComBat was processed using Numerical Python (NumPy) (version 1.23.5) and Pandas (version 1.5.3) [29], [30]. Afterward, Least Absolute Shrinkage and Selection Operator with Cross Validated (LassoCV) and Recursive Feature Elimination with Cross-Validated (RFECV) from Scikit-learn (version 1.2.2) were implemented [31]. Data scaling was done using StandardScaler() function. For data visualization, time estimation and model handling, Python libraries such as Matplotlib (version 3.7.1), Seaborn (version 0.12.2), Time (version 1.23.5) and Joblib (version 1.3.2)were used [32]–[35].

At first, LassoCV was applied with five times cross validation. LassoCV determined the optimum parameters for Lasso algorithm regularization. Lasso algorithm reduced the sum of the absolute values of the regression coefficients which reduced overfitting. LassoCV determined the important features to simplify the model, which was suitable for high-dimensional data analysis. Secondly, RFECV, with five times cross validations, chose the features by continually deleting the least significant features and evaluating the performance of the model after each reduction. This lessened the possibility of overfitting and aided in identifying the most important features.

### Training of machine learning based classifier models

The most important features were applied to train the classification models of Scikit-learn. To do so, the dataset was splitted in two sections called train and test datasets. Seventy percent data was implemented for training and thirty percent data was used for testing each classifier. Classifiers such as logistic regression, random forest, naive bayes classifier, k-nearest neighbors (parameters: n_neighbors=11, metric=’manhattan’, weights=’uniform’), and support vector machine were trained and tested. Moreover, a deep neural network based classifier was built using Tensorflow (version: 2.12.0) [36], [37]. All models went through ten times cross validation with shuffled train and test datasets. The performance of the models were evaluated with Precision, Recall, F1-Score, Mean accuracy, Matthews correlation coefficient (MCC) score, Area under the Receiver operating characteristic curve (AUC-ROC) and Confusion matrix.

## Result

### DEG Analysis Showed Profound Gene Expression Differences in Blood Samples from PD Patient

In PD patient samples, a sum of 13,248 genes exhibiting differential expression (DEGs) was detected. Among these, 7630 genes demonstrated upregulation, while 5618 genes displayed downregulation. Subsequently, applying an adjusted p-value threshold (<0.05) narrowed down the gene list to 3,980 genes. Further refinement involved filtering genes by their median expression levels, preserving those with values equal to or higher than the calculated median (1,990 genes). Lastly, a final subset of genes was derived by additional filtration using specific criteria on average expression levels (AveExpr > 3). This stringent process resulted in the identification of 678 DEGs lists of which along with their Log2 FC values, average Log2 FCs are provided in Supplementary Table 1. The R Studio and library tidyverse package were employed to create a volcano plot for contrasting the differentially expressed genes (DEGs) between PD patients and healthy controls. The plot in the Figure 3a showcased ten noteworthy DEGs (*RND1, COG8 /// PDF, GPR107, FBXW4P1, PPP2R2D, F7, OAZ1, MYL12A, TSPYL2, ANKRD36B*) where upregulated DEGs were represented on the right side in yellow, and downregulated DEGs appeared on the left side in red.

**Figure 3:**
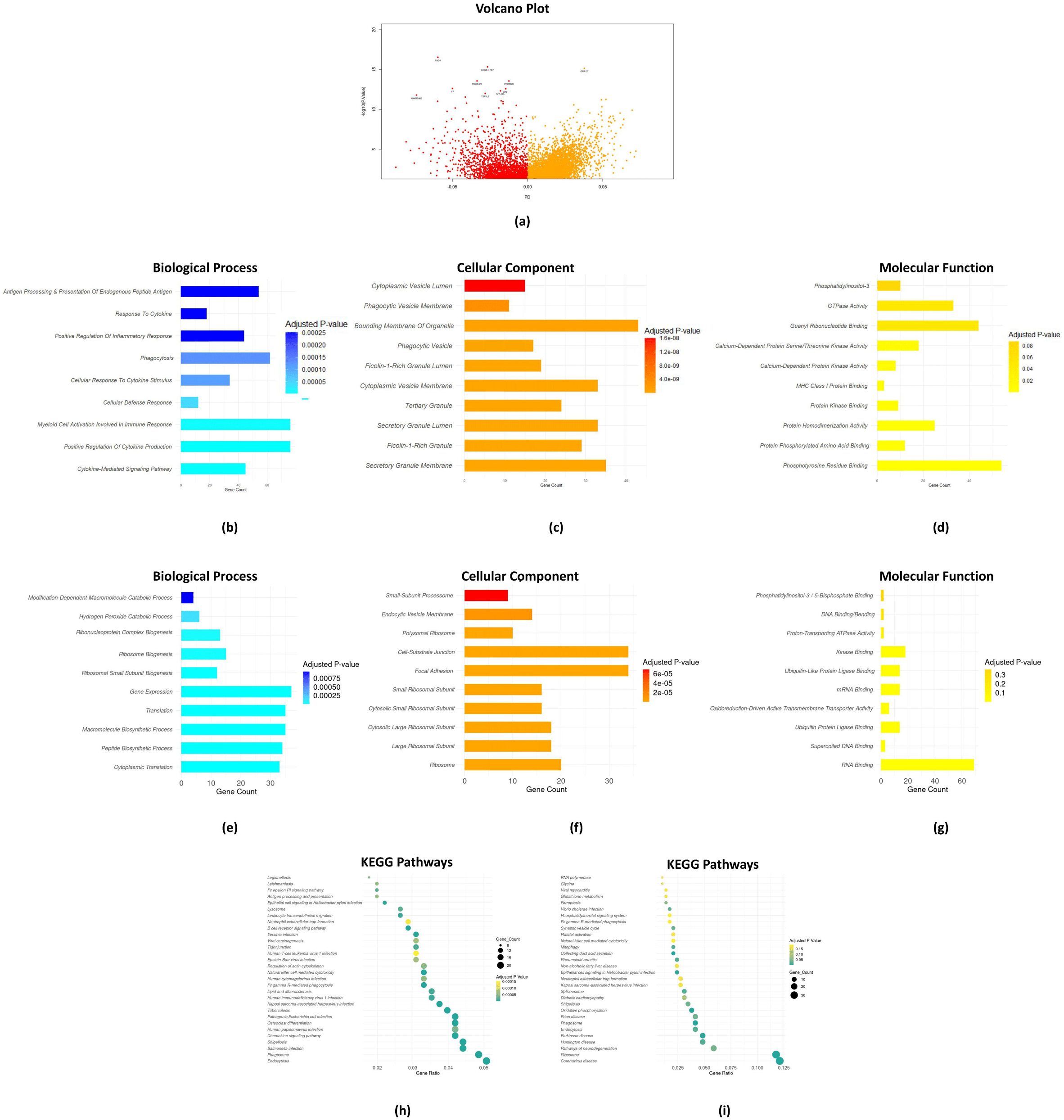
Identification and functional annotation of DEGs. a) A volcano plot of the distribution of all DEGs in Parkinson’s disease,, mapping the 7630 upregulated genes (yellow region) and 5618 downregulated genes (red region); b) GO analysis of the upregulated DEGs in BP terms; c) GO analysis of the upregulated DEGs in CC terms; d) GO analysis of the upregulated DEGs in MF terms; e) GO analysis of the downregulated DEGs in BP terms; f) GO analysis of the downregulated DEGs in CC terms; g) GO analysis of the downregulated DEGs in MF terms; The ordinate represents the GO term name, and the abscissa shows the count of DEGs enriched in the GO term. The color of the bar represents the p-value associated with the enriched term. KEGG pathway enrichment of the h) upregulated DEGs and (D) downregulated DEGs. The ordinate represents the name of the KEGG pathway, and the abscissa shows the ratio of enriched gene counts within the pathways.The count of DEGs is represented by bubble size and P-value by color. DEGs differentially expressed genes; GO, Gene Ontology; BP Biological Process; CC Cellular component; MF Molecular Function; KEGG, Kyoto Encyclopedia of Genes and Genomes.

### Functional and Pathway Enrichment Analysis Revealed the Underlying Biology of PD

To further understand the functional roles and potential pathways involved in the identified DEGs, functional annotation and pathway analysis, including GO and KEGG, were performed. For this purpose, only GO terms displaying significant relevance with a P-value < 0.05 were considered (Supplementary Table 2). The most significant ten gene-enriched GO terms, both for down-regulated and up-regulated genes, were depicted in Figure 3. Focusing on the upregulated DEGs, the GO term enrichment analysis unveiled a diverse array of associations across cellular components, biological processes, and molecular functions. Specifically, the outcomes of the GO analysis were distinctly indicative of significant enrichment in various biological processes, ranging from responses to the cytokine-mediated signaling pathway, positive regulation of cytokine production, myeloid cell activation involved in immune response, to the antigen processing and presentation of endogenous peptide antigen via MHC Class I Via ER Pathway, among several others (Figure3b). In terms of cellular components, most of the genes were localized to the bounding membrane of organelle, secretory granules, and cytoplasmic vesicle membrane (Figure3c). Moreover, molecular functions linked to phosphotyrosine residue binding, protein phosphorylated amino acid and protein kinase binding and protein homodimerization activity were notably enriched (in terms of p-value). Only a few genes were attributed to molecular functions like calcium-dependent protein kinase activity and MHC class I protein binding (Figure 3d).

In contrast, among the downregulated DEGs, the functional landscape revealed a distinct profile. In the context of biological processes within the cell, they were significantly involved in activities like gene expression, cytoplasmic translation, and various biosynthetic processes. Notably, fewer genes were linked to catabolic processes, such as breaking down hydrogen peroxide or modification-dependent large molecules (Figure 3e). Regarding cellular components, they were frequently found in cell parts like cell-substrate junctions, focal adhesions, ribosomes and small and large ribosomal and cytosolic ribosomal subunits (Figure 3f). Additionally, these genes were mainly linked to RNA binding in terms of molecular functions (Figure 3g).

To further investigate the biological functions of differentially expressed genes, we performed enrichment analyses by mapping the sequences to the KEGG database categories. For upregulated DEGs, our results indicated that 337 upregulating DEGs were assigned to 135 KEGG pathways in total. The pathway with the most DEGs was ‘Endocytosis’, followed by ‘Phagosome’, ‘Shigellosis’ and ‘*Salmonella* infection’. According to the adjusted p-value for the pathway enrichment analysis, most DEGs were significantly enriched in “Cellular Homeostasis and Signaling Pathways in Immune Response.”, such as Phagosome, Osteoclast differentiation, Endocytosis, Fc gamma R-mediated phagocytosis and Chemokine signaling pathway (Figure 3h).

Regarding the downregulated DEGs, the results exhibited an association of 341 genes with a total of 236 KEGG pathways. Notably, certain genes exhibited significant enrichment in widely recognized pathways related to PD, such ribosome, phagosome, endocytosis, and pathways of neurodegeneration as depicted in the Figure 3i.

### *RPS28* downregulation and *HLA-F, IRF1* upregulation is critical in PD

The analysis of protein-protein interactions (PPI) for the upregulated DEGs highlighted a rich network with a total of 326 proteins (nodes) and 1671 interactions (edges). The ten most interconnected DEGs included genes like *MAPK3*, *RAC1*, *GRB2*, *IRF2*, *HLA-F*, *HLA-B*, *HLA-C*, *IRF7*, *ADAR*, and *IRF1* depicted in vibrant yellow tones in the Figure 4a. Interestingly, all of these hub molecules have previously been associated with PD [38]–[43]. Nevertheless, two genes, *HLA-F* and *IRF1*, exhibited the most noteworthy statistical significance in relation to PD with a p-value ≤ 0.001 (Figure 5a).

**Figure 4:**
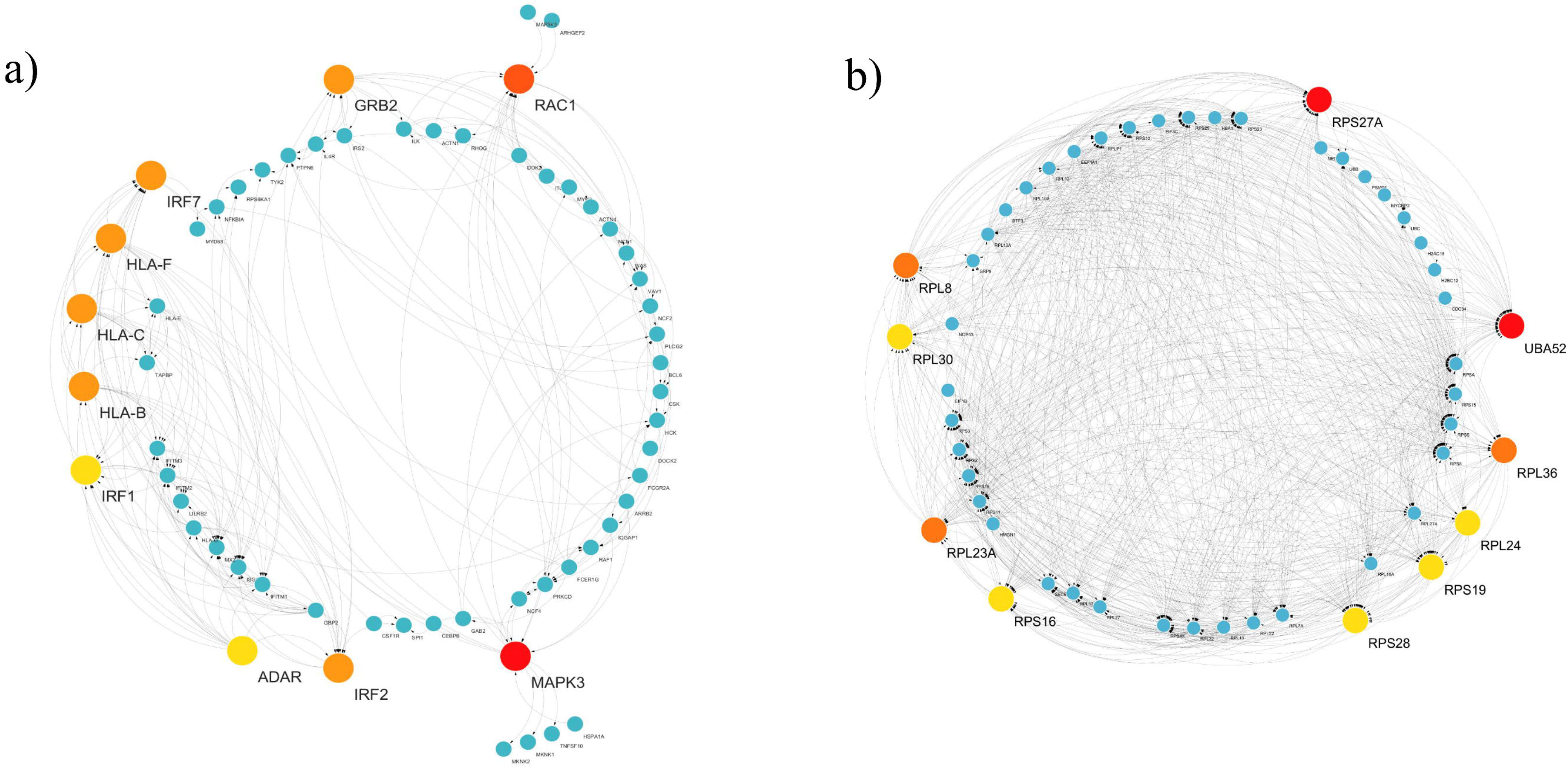
PPI networks of differentially expressed genes (DEGs). a) PPI network for DEGs from upregulated genes; b) PPI network for DEGs from downregulated genes. Highly connected proteins are highlighted in red, with node size reflecting the degree of their connectivity.

**Figure 5:**
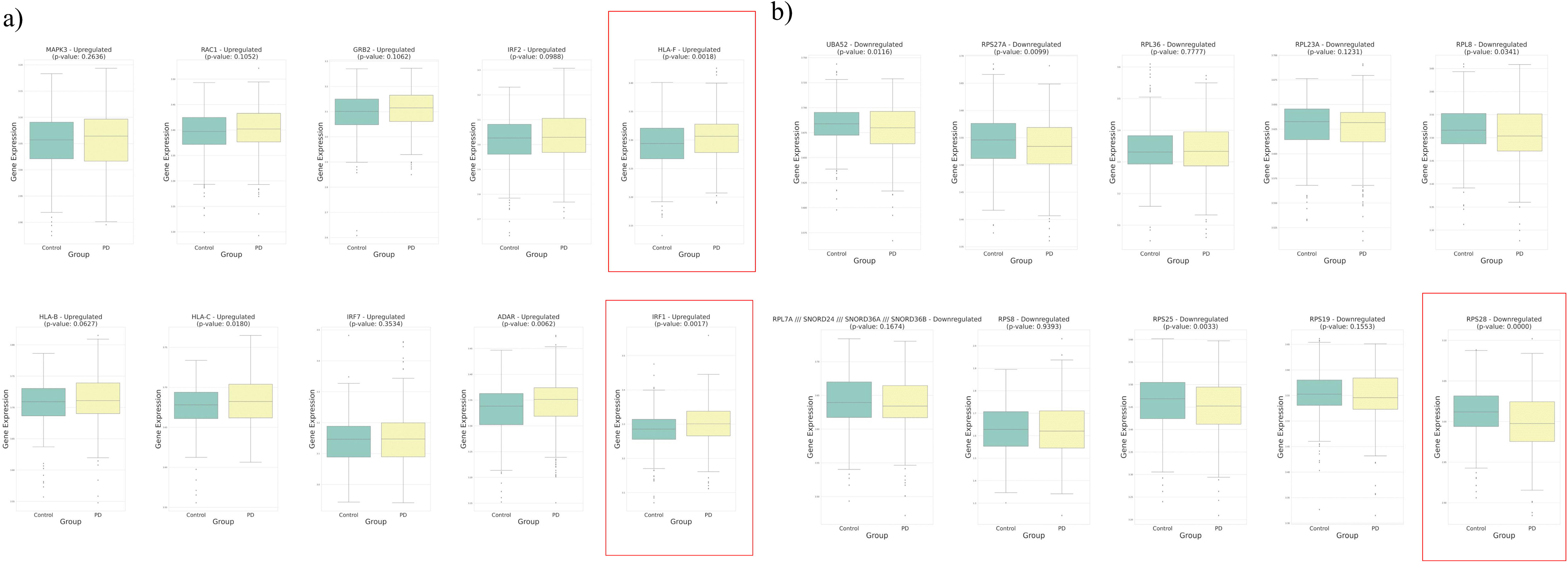
Boxplot depicting the top 20 highly connected Differentially Expressed Genes identified from the Protein-Protein Interaction (PPI) network analysis. Subsets of these genes include: a) 10 hub DEGs from upregulated genes; b) 10 hub DEGs from downregulated genes. Statistically significant differences, as determined by an Independent T-test, are highlighted with red boxes (p ≤ 0.001), emphasizing their distinct expression profiles associated with Parkinson’s disease.

Similarly, the PPI network of downregulated DEGs revealed an intricate interaction among 310 nodes and 2129 edges. The top ten enriched DEGs here are *UBA52*, *RPS27A*, *RPL36*, *RPL23A*, *RPL8*, *RPL7A*, *RPS8*, *RPS25*, *RPS19*, and *RPS28* with the most highly connected proteins highlighted in red (Figure 4b). Among them *RPS28* demonstrated statistically significant associations with PD (p value ≤ 0.001) (Figure 5b) [44]–[46] .

### Support Vector machine Achieved Optimal Classification Performance

Batch corrected data had 13249 genes (features). LassoCV discarded 13205 features and selected 44 genes. REFCV discarded 1 gene and finally 43 genes were implemented to train the machine learning models. The selected genes/ features were ’*ABCA5’, ’ANXA3’, ’ARG1’, ’ARL17A /// ARL17B’, ’BTN3A2’, ’CASC1’, ’COL9A3’, ’DEFA4’, ’DRD4’, ’DSP’, ’ENC1’, ’FCGR1A’, ’FCGR1A /// FCGR1B /// FCGR1C’, ’FKBP14’, ’FOXD1’, ’FPGT’, ’GPX3’,’GUSBP3 /// SMA4 /// SMA5’, ’GYS2’, ’HLA-DQA1’, ’HS3ST3A1’, ’IGJ’, ’IL18R1’, ’KIR2DL3’, ’KRT1’, ’LARP6’, ’LOC644172 /// MAPK8IP1’, ’LRRN3’, ’MCTP1’, ’MYH7B’, ’NMBR’, ’PCF11’, ’PF4V1’, ’POMZP3’, ’PTGDS’, ’RPS4Y1’, ’SCD5’, ’SLFN12’, ’TJP2’, ’TUBB2A’, ’XIST’, ’XK*’. Classifiers models of Scikit-learn such as logistic regression, random forest, naive bayes classifier, k-nearest neighbors and support vector machine were trained and tested using these 43 genes (Supplementary Figure 1). Moreover, using Tensorflow, an Artificial Neural Network (ANN) with 128 neurons in the first layer, 64 neurons in the second layer and 2 neurons in the last layer was constructed. The activation function for the first and second layer were relu. In the third layer, softmax activation function was given. Models performances were evaluated with Precision, Recall, F1-Score, Mean accuracy and Area under the Receiver operating characteristic curve (AUC-ROC). The Precision values of the models were 0.6-0.7. Precision means the percentage of positive instances that are actually positive. Recall value was observed in between 0.46-0.80. Recall determines the percentage of instances that are actually positive and are expected to be positive.

The range of F1 score value was 0.55-0.68. The F1-score is calculated as an average of precision and recall. MCC value was observed from 0.2-0.35. Using all four values of the confusion matrix, MCC assesses the robustness of binary classifications that are more reliable than precision, recall, and accuracy. The AUC-ROC was between 0.66-0.71 . The capacity of a binary classifier to distinguish between positive and negative instances is measured by the AUC-ROC. The range of mean accuracy was 0.6-0.64. Accuracy measures the proportion of all instances that are correctly classified. The time that was taken to train the models was 0.01-30 seconds. ANN took the highest time where naive bayes classifier was the fastest model to train.

## Discussion

Parkinson’s disease (PD) is a complex neurodegenerative disorder with various genetic and environmental factors contributing to its development [47]. Currently, there is no cure for PD, and slowing down its progression remains a challenge [11]. The neurodegenerative process in PD begins long before any visible symptoms appear. Thus there remains a pressing need of the identification of biomarkers that could aid in the diagnosis and treatment of PD, especially in its early stages when neuroprotective therapies may be more effective. Discovering PD biomarkers could be highly beneficial for identifying individuals at risk, diagnosing the disease, and monitoring its progression. In light of this, blood biomarkers that are relatively easy to obtain would have a great potential for use if they are able to meet the necessary requirements in a research setting and clinical practice.

The underlying factors leading to neuroinflammation and subsequent dopaminergic neuron loss in the substantia nigra during PD progression are already well-established [48]. The build-up of inflammatory substances and the activation of microglia in affected brain areas are significant contributors to this phenomenon, consistent with the GO analysis of our study’s Differentially Expressed Genes (DEGs) (Figure 3). A study conducted by Iannaccone et al., identified activated microglia in the brains of PD patients, initiating a cycle of inflammation and neuronal harm that ultimately results in the degeneration of dopamine neurons in PD [48]. However, in addition to resident immune cells like microglia, the involvement of peripheral immune cells, specifically blood-borne myeloid cells, has emerged as a critical facet in PD susceptibility and progression [49]. Moreover, dysregulated cytokines in the blood, cerebrospinal fluid, and brain of PD patients suggest a connection between peripheral and central immune systems [50]. Quantitative and qualitative changes in leukocytes (which are the immune cells in peripheral blood) and their subpopulations have also been observed in PD patients. [51], [52]. Besides, blood cells also get affected by PD in different ways. A recent study reported that PD patients have higher blood pressure variability [53]. This variability might affect the vital processes such as blood flow and oxygen amount that will eventually affect the gene expression pattern of the PD patients. Correspondingly, aging and neuroinflammation can disrupt the blood-brain barrier (BBB) and cause neurovascular problems in PD, potentially involving peripheral cells in neurodegeneration [54]. The BBB, along with other protective barriers, shields brain neurons from external threats, and changes in the BBB associated with aging are a significant risk factor for PD [55]. Inflammatory markers like cytokines and chemokines in the peripheral system that play a key role in immune regulation, cross the BBB and impact the central nervous system, Therefore, peripheral inflammation could be a significant contributor to the etiology of PD as well as to disease progression [52], [55].

The PPI network analysis of upregulated genes highlighted key hub proteins related to immune and inflammatory processes, consistent with Gene Ontology results. Notably, *HLA-F*, a class I histocompatibility antigen and *IRF1* (interferon regulatory factor 1), a transcriptional regulator and tumor suppressor activating immune-related genes, were identified as most statistically significant genes (Figure 5a) [56], [57]. While the association between HLA class I and Parkinson’s disease dates back over four decades, recent research has underscored a more clearer and more replicable role of HLA class II in PD predisposition and protection [41], [58], [59]. Therefore, further exploration is needed to understand the potential association of HLA-class I proteins, specifically *HLA-F*, with PD. Additionally, IRFs have a broader and more direct impact on inflammation and immunity by controlling cell development and function, which are essential for the immune response [60]. In a recent study using a mouse model of PD, *IRF-1* was found to be upregulated resulting from MDM2-mediated ubiquitylation through the ubiquitin proteasome pathway[61]. These findings suggest *HLA-F* and *IRF1* as potential biomarkers, offering insights into their roles in immune responses and Parkinson’s disease pathogenesis.

In parallel, among the proteins with the highest number of interactions in the downregulated genes network, we identified *RPS28* as the most statistically significant DEG, as depicted in the figure 5b. Ribosomal protein S28 (*RPS28*) gene encodes a ribosomal protein that is a component of the 40S subunit [62]. Ribosomal proteins play a crucial in the intricate process of cellular protein synthesis, including initiation and elongation. These proteins also possess the unique ability to regulate their own synthesis at the translational level [63], [64]. Intriguingly, a study investigating Parkinson’s disease progression in Braak and middle-aged individuals unveiled a connection between alterations in the expression of ribosomal protein genes and the advancement of the disease. This relationship exhibits a level of complexity that not only varies across different disease stages but also among specific regions of the brain [65]. Adding to our study, the down-regulation of *RPS28* in PD had been observed in a similar study using Mass Spectrometry–Based Proteomics Analysis of Human Substantia Nigra [66]. Taken together, our analyses indicate that *HLA-F*, *IRF1*, and *RPS28* could potentially serve as valuable markers for identifying Parkinson’s disease patients at a stage when therapeutic interventions could be most effective, as well as for monitoring the severity of the disease. Furthermore, the results can be supported by machine learning based classifiers for two step validation.

To develop a machine learning based prediction model, we trained classifiers such as logistic regression, support vector machine, k-nearest neighbor, naive bayes classifier, random forest and artificial neural network with 568 samples. Here, 273 samples (∼ 48%) were from the healthy population and 295 (∼ 52%) from the PD population where each sample had data points from 13249 genes. The whole dataset went through log2 transformation and an empirical Bayes framework to adjust the batch effect. Of 13249 genes, LassoCV followed by REFCV only suggested 43 genes for model training. Among all the models, the Support Vector Machine (SVM) has the most optimum performance based on AUC-ROC, MCC and Accuracy (Figure 6). SVM showed ∼65% accuracy (95%CI: 0.58-0.66), 0.70 AUC-ROC (95%CI: 0.70-0.71), 0.35 MCC (95%CI:0.34 - 0.39). SVM took only 0.115 seconds for model training. The model has better F1 score, precision and recall values (Figure 6). Hence, SVM had the best capability to distinguish PD and healthy populations from blood based gene expressions. However, it showed 16.96% false negative results which is higher than k-nearest neighbor (9.94%) (Figure 7). On the other hand, naive bayes classifier and logistic regression showed lowest false positive (15.79%) results, however, these models had higher false negative outcomes with lower MCC scores (Figure 7). Finally, we emphsesized SVM since MCC score is a more effective metric for binary classifications in biological domains [67], [68].

**Figure 6:**
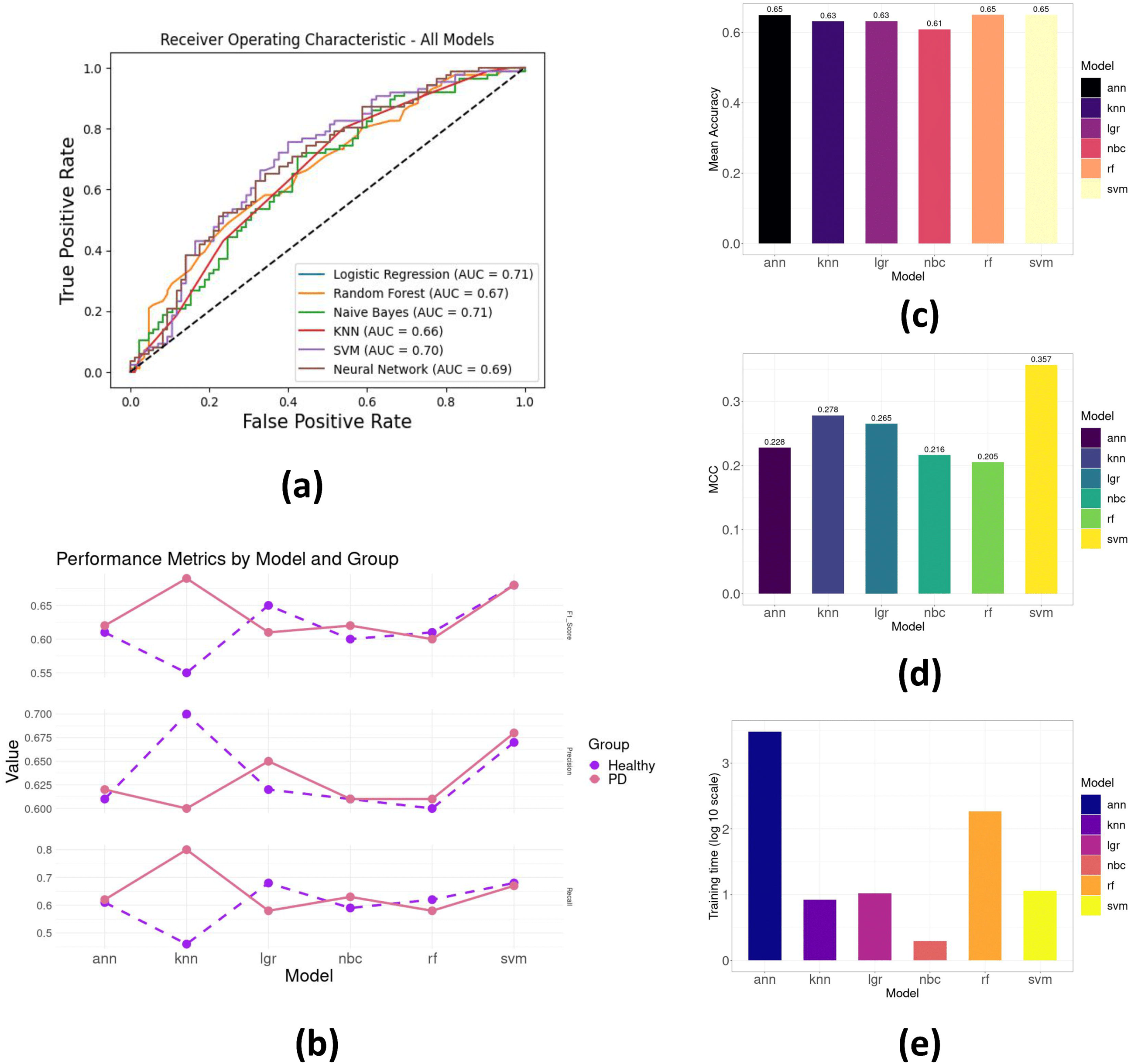
Evaluation results of the machine learning models: a) Area under the ROC Curve (AUC-ROC); b) F1 score, Precision and Recall (the plain lines represent the results for PD detection and the dashed lines represent the results from healthy population); c) Mean accuracy (Mean of 10 fold cross validation results); d) Matthews Correlation Coefficient (MCC) score and e) Training time required for the models where the time has been demonstrated in log 10 scale.

**Figure 7:**
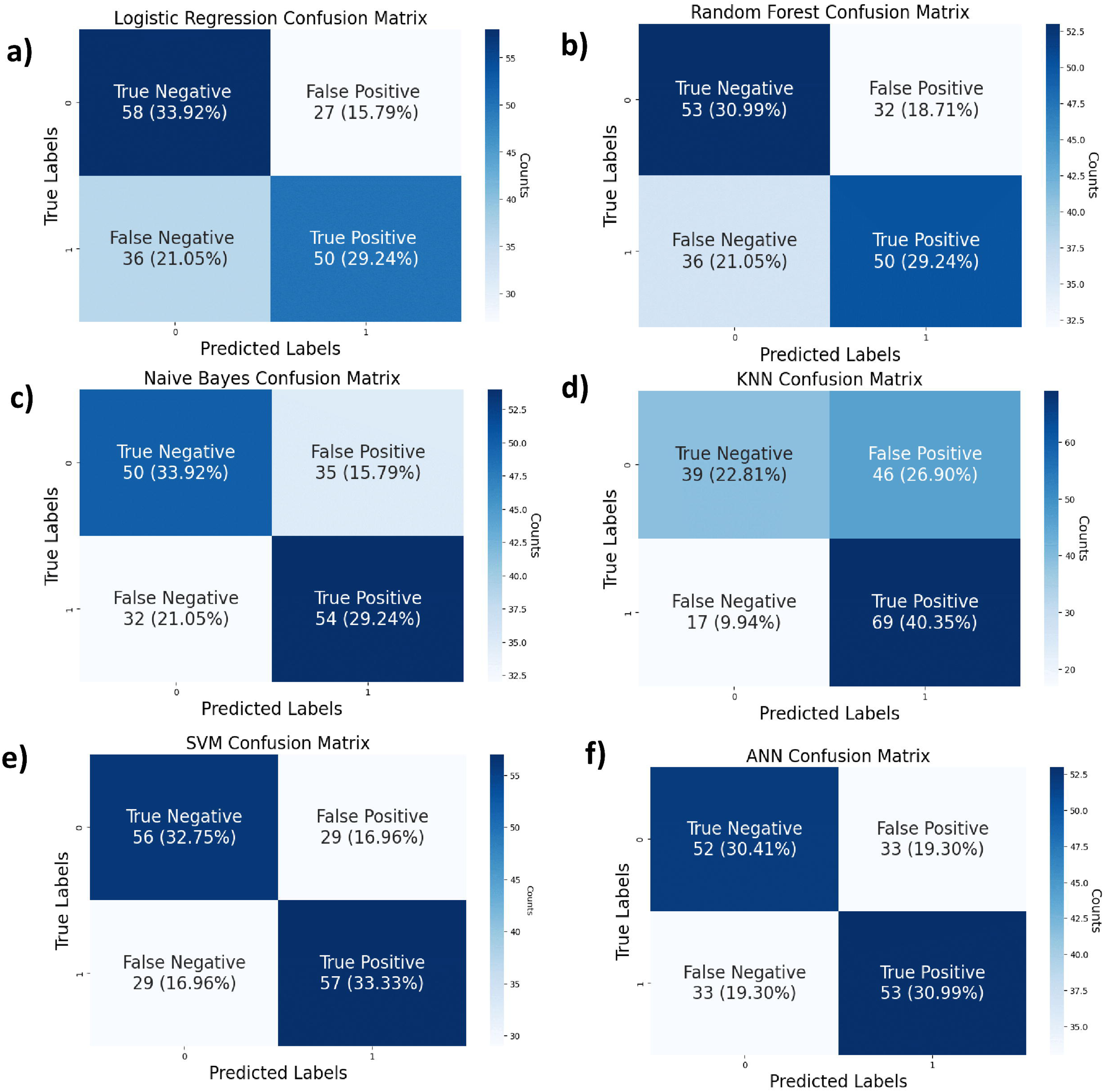
Confusion matrix of the trained classifiers. Here, the True Positive (TP), True Negative (TN), False positive (FP) and False Negative (FN) values of a) Logistic Regression, b) Random Forest, c) Naive Bayes Classifier, d) K-Nearest Neighbors (KNN), e) Support Vector Machine (SVM) and f) Artificial Neural Networks (ANN) have been depicted. The dark blue gradient resembles higher values whereas light blue gradient represents lower values in the matrix tables.

The results imply that for each 100 individuals, around 15 to 16 healthy people will be diagnosed with PD and ∼ 17 PD patients will be misdiagnosed as a healthy person. Yet, ∼67 individuals will receive an accurate diagnosis using this non-invasive technique. Therefore, this could be used for a preliminary screening for PD, which would significantly reduce costs and sufferings regarding PD diagnosis. In future, more gene expression data from PD patients, especially from non-european countries, will improve the performance of this model. The trained model was deposited in Github [69]. The repository name for this model is **P**ark**I**nson’s disease detection **T**ool using blood-based micro**Array** data (PitArray) [70]. From this repository, the model can be used via Google Colab or local computer [71].

### Conclusion

The increasing prevalence of Parkinson’s disease is a global health concern, and while our understanding of its causes has improved, diagnosis remains challenging. Thus, this research aims to advance non-invasive diagnostic methods for PD through molecular analysis and machine learning approaches.

## Author Contributions

A.B., T.B.J., A.B.L. performed data collection, preprocessing, differential gene expression analysis and machine learning. A.B., I.A., M.S.A., and T.B.J. developed the PitArray tool. A.B. and T.B.J wrote the original draft of the manuscript. Z.M.C., M.U.H., K.C.D., and C.A.K. reviewed and edited the manuscript. M.S. supervised the study.

## Data Availability

The data generated in this study are included within the manuscript and the supplementary files.

## Code Availability

Code for using the PitArray tool can be found at: https://github.com/Arittra95/PitArray.

## Author Biographies

**Arittra Bhattacharjee**

Arittra Bhattacharjee is a Scientific Officer at the Bioinformatics Division, National Institute of Biotechnology.

**Tabassum Binte Jamal**

Tabassum Binte Jamal is a Post-Graduate Research Fellow at the Bioinformatics Division, National Institute of Biotechnology.

**Ishtiaque Ahammad**

Ishtiaque Ahammad is a Scientific Officer at the Bioinformatics Division, National Institute of Biotechnology.

**Anika Bushra Lamisa**

Anika Bushra Lamisa is a Post-Graduate Research Fellow at the Bioinformatics Division, National Institute of Biotechnology.

**Md. Shamsul Arefin**

Md. Shamsul Arefin is a graduate student at the Department of Biochemistry and Microbiology, North South University.

**Zeshan Mahmud Chowdhury**

Zeshan Mahmud Chowdhury is a Scientific Officer at the Bioinformatics Division, National Institute of Biotechnology.

**Mohammad Uzzal Hossain**

Mohammad Uzzal Hossain is a Senior Scientific Officer (C.C.) at the Bioinformatics Division, National Institute of Biotechnology.

**Keshob Chandra Das**

Keshob Chandra Das is a Chief Scientific Officer (C.C.) at the Molecular Biotechnology Division, National Institute of Biotechnology.

**Dr. Chaman Ara Keya**

Dr. Chaman Ara Keya is an Associate Professor at the Department of Biochemistry and Microbiology, North South University.

**Dr. Md Salimullah**

Dr. Md Salimullah is the Director General (A.C.) and Chief Scientific Officer at the National Institute of Biotechnology.

## Supplementary Files

**Supplementary Figure 1:** The log2-fold (in log10 scale) value of the up-regulated genes (Red lollipop bars) and downregulated genes (Blue lollipop bars) that were used as features for machine learning. Fifteen genes were down regulated and 27 genes were up-regulated (total 42 genes).

**Supplementary Table 1:** List of 678 upregulated and downregulated differentially expressed genes.

**Supplementary Table 2:** Top 10 statistically significant GO and KEGG pathway enrichment analyses for the upregulated and downregulated differentially expressed genes. GO, Gene Ontology; KEGG, Kyoto Encyclopedia of Genes and Genomes.

## References

[1] W. Poewe et al., “Parkinson disease,” Nat. Rev. Dis. Primer, vol. 3, no. 1, p. 17013, Mar. 2017, doi: 10.1038/nrdp.2017.13.

[2] “Parkinson disease.” https://www.who.int/news-room/fact-sheets/detail/parkinson-disease (accessed Sep. 19, 2023).

[3] “Parkinson’s Disease: Challenges, Progress, and Promise,” National Institute of Neurological Disorders and Stroke. https://www.ninds.nih.gov/current-research/focus-disorders/focus-parkinsons-disease-research/parkinsons-disease-challenges-progress-and-promise (accessed Sep. 19, 2023).

[4] A. Kouli, K. M. Torsney, and W.-L. Kuan, “Parkinson’s Disease: Etiology, Neuropathology, and Pathogenesis,” in Parkinson’s Disease: Pathogenesis and Clinical Aspects, T. B. Stoker and J. C. Greenland, Eds., Brisbane (AU): Codon Publications, 2018. Accessed: Sep. 19, 2023. [Online]. Available: http://www.ncbi.nlm.nih.gov/books/NBK536722/

[5] D. W. Dickson et al., “Neuropathological assessment of Parkinson’s disease: refining the diagnostic criteria,” Lancet Neurol., vol. 8, no. 12, pp. 1150–1157, Dec. 2009, doi: 10.1016/S1474-4422(09)70238-8.

[6] P. Damier, E. C. Hirsch, Y. Agid, and A. M. Graybiel, “The substantia nigra of the human brain,” Brain, vol. 122, no. 8, pp. 1437–1448, Aug. 1999, doi: 10.1093/brain/122.8.1437.

[7] D. Iacono et al., “Parkinson disease and incidental Lewy body disease: Just a question of time?,” Neurology, vol. 85, no. 19, pp. 1670–1679, Nov. 2015, doi: 10.1212/WNL.0000000000002102.

[8] S. Szatmari, B. Min-Woo Illigens, T. Siepmann, A. Pinter, A. Takats, and D. Bereczki, “Neuropsychiatric symptoms in untreated Parkinson’s disease,” Neuropsychiatr. Dis. Treat., vol. Volume 13, pp. 815–826, Mar. 2017, doi: 10.2147/NDT.S130997.

[9] L. M. Zuckerman, “Parkinson_s Disease and the Orthopaedic Patient:,” J. Am. Acad. Orthop. Surg., vol. 17, no. 1, pp. 48–55, Jan. 2009, doi: 10.5435/00124635-200901000-00007.

[10] M. J. Armstrong and M. S. Okun, “Diagnosis and Treatment of Parkinson Disease: A Review,” JAMA, vol. 323, no. 6, p. 548, Feb. 2020, doi: 10.1001/jama.2019.22360.

[11] “Parkinson disease.” https://www.who.int/news-room/fact-sheets/detail/parkinson-disease (accessed Sep. 19, 2023).

[12] M. Agrawal and A. Biswas, “Molecular diagnostics of neurodegenerative disorders,” Front. Mol. Biosci., vol. 2, Sep. 2015, doi: 10.3389/fmolb.2015.00054.

[13] M. Funayama, K. Nishioka, Y. Li, and N. Hattori, “Molecular genetics of Parkinson’s disease: Contributions and global trends,” J. Hum. Genet., vol. 68, no. 3, pp. 125–130, Mar. 2023, doi: 10.1038/s10038-022-01058-5.

[14] E. Mariani, F. Frabetti, A. Tarozzi, M. C. Pelleri, F. Pizzetti, and R. Casadei, “Meta-Analysis of Parkinson’s Disease Transcriptome Data Using TRAM Software: Whole Substantia Nigra Tissue and Single Dopamine Neuron Differential Gene Expression,” PLOS ONE, vol. 11, no. 9, p. e0161567, Sep. 2016, doi: 10.1371/journal.pone.0161567.

[15] L. Su, C. Wang, C. Zheng, H. Wei, and X. Song, “A meta-analysis of public microarray data identifies biological regulatory networks in Parkinson’s disease,” BMC Med. Genomics, vol. 11, no. 1, p. 40, Dec. 2018, doi: 10.1186/s12920-018-0357-7.

[16] D. M. Phung, J. Lee, S. Hong, Y. E. Kim, J. Yoon, and Y. J. Kim, “Meta-Analysis of Differentially Expressed Genes in the Substantia Nigra in Parkinson’s Disease Supports Phenotype-Specific Transcriptome Changes,” Front. Neurosci., vol. 14, p. 596105, Dec. 2020, doi: 10.3389/fnins.2020.596105.

[17] M. Falchetti, R. D. Prediger, and A. Zanotto-Filho, “Classification algorithms applied to blood-based transcriptome meta-analysis to predict idiopathic Parkinson’s disease,” Comput. Biol. Med., vol. 124, p. 103925, Sep. 2020, doi: 10.1016/j.compbiomed.2020.103925.

[18] J. Augustine and A. S. Jereesh, “Blood-based gene-expression biomarkers identification for the non-invasive diagnosis of Parkinson’s disease using two-layer hybrid feature selection,” Gene, vol. 823, p. 146366, May 2022, doi: 10.1016/j.gene.2022.146366.

[19] E. A. Welsh, S. A. Eschrich, A. E. Berglund, and D. A. Fenstermacher, “Iterative rank-order normalization of gene expression microarray data,” BMC Bioinformatics, vol. 14, no. 1, p. 153, May 2013, doi: 10.1186/1471-2105-14-153.

[20] P. Kupfer, R. Guthke, D. Pohlers, R. Huber, D. Koczan, and R. W. Kinne, “Batch correction of microarray data substantially improves the identification of genes differentially expressed in Rheumatoid Arthritis and Osteoarthritis,” BMC Med. Genomics, vol. 5, no. 1, p. 23, Dec. 2012, doi: 10.1186/1755-8794-5-23.

[21] C. Chen et al., “Removing Batch Effects in Analysis of Expression Microarray Data: An Evaluation of Six Batch Adjustment Methods,” PLoS ONE, vol. 6, no. 2, p. e17238, Feb. 2011, doi: 10.1371/journal.pone.0017238.

[22] M. Ashburner et al., “Gene Ontology: tool for the unification of biology,” Nat. Genet., vol. 25, no. 1, pp. 25–29, May 2000, doi: 10.1038/75556.

[23] A. Tomczak et al., “Interpretation of biological experiments changes with evolution of the Gene Ontology and its annotations,” Sci. Rep., vol. 8, no. 1, p. 5115, Mar. 2018, doi: 10.1038/s41598-018-23395-2.

[24] M. Kanehisa, “KEGG: Kyoto Encyclopedia of Genes and Genomes,” Nucleic Acids Res., vol. 28, no. 1, pp. 27–30, Jan. 2000, doi: 10.1093/nar/28.1.27.

[25] M. V. Kuleshov et al., “Enrichr: a comprehensive gene set enrichment analysis web server 2016 update,” Nucleic Acids Res., vol. 44, no. W1, pp. W90–W97, Jul. 2016, doi: 10.1093/nar/gkw377.

[26] E. Y. Chen et al., “Enrichr: interactive and collaborative HTML5 gene list enrichment analysis tool,” BMC Bioinformatics, vol. 14, no. 1, p. 128, Dec. 2013, doi: 10.1186/1471-2105-14-128.

[27] “About - STRING functional protein association networks.” https://string-db.org/cgi/about (accessed Sep. 19, 2023).

[28] “Cytoscape App Store - cytoHubba.” https://apps.cytoscape.org/apps/cytohubba (accessed Sep. 19, 2023).

[29] C. R. Harris et al., “Array programming with NumPy,” Nature, vol. 585, no. 7825, pp. 357–362, Sep. 2020, doi: 10.1038/s41586-020-2649-2.

[30] W. McKinney, “Data Structures for Statistical Computing in Python,” presented at the Python in Science Conference, Austin, Texas, 2010, pp. 56–61. doi: 10.25080/Majora-92bf1922-00a.

[31] F. Pedregosa et al., “Scikit-learn: Machine Learning in Python,” J. Mach. Learn. Res., vol. 12, no. 85, pp. 2825–2830, 2011.

[32] “Joblib: running Python functions as pipeline jobs — joblib 1.3.2 documentation.” https://joblib.readthedocs.io/en/stable/ (accessed Sep. 19, 2023).

[33] J. D. Hunter, “Matplotlib: A 2D Graphics Environment,” Comput. Sci. Eng., vol. 9, no. 3, pp. 90–95, 2007, doi: 10.1109/MCSE.2007.55.

[34] M. Waskom, “seaborn: statistical data visualization,” J. Open Source Softw., vol. 6, no. 60, p. 3021, Apr. 2021, doi: 10.21105/joss.03021.

[35] “time — Time access and conversions,” Python documentation. https://docs.python.org/3/library/time.html (accessed Sep. 19, 2023).

[36] M. Abadi et al., “TensorFlow: a system for large-scale machine learning,” in Proceedings of the 12th USENIX conference on Operating Systems Design and Implementation, in OSDI’16. USA: USENIX Association, Nov. 2016, pp. 265–283.

[37] V. B. Mathema, P. Sen, S. Lamichhane, M. Orešič, and S. Khoomrung, “Deep learning facilitates multi-data type analysis and predictive biomarker discovery in cancer precision medicine,” Comput. Struct. Biotechnol. J., vol. 21, pp. 1372–1382, 2023, doi: 10.1016/j.csbj.2023.01.043.

[38] A. Bohush, G. Niewiadomska, and A. Filipek, “Role of Mitogen Activated Protein Kinase Signaling in Parkinson’s Disease,” Int. J. Mol. Sci., vol. 19, no. 10, p. 2973, Sep. 2018, doi: 10.3390/ijms19102973.

[39] H. Kim et al., “The Small GTPase RAC1/CED-10 Is Essential in Maintaining Dopaminergic Neuron Function and Survival Against α-Synuclein-Induced Toxicity,” Mol. Neurobiol., vol. 55, no. 9, pp. 7533–7552, Sep. 2018, doi: 10.1007/s12035-018-0881-7.

[40] K. Lei et al., “Immune-associated biomarkers for early diagnosis of Parkinson’s disease based on hematological lncRNA–mRNA co-expression,” Biosci. Rep., vol. 40, no. 12, p. BSR20202921, Dec. 2020, doi: 10.1042/BSR20202921.

[41] A. Puschmann et al., “Human leukocyte antigen variation and Parkinson’s disease,” Parkinsonism Relat. Disord., vol. 17, no. 5, pp. 376–378, Jun. 2011, doi: 10.1016/j.parkreldis.2011.03.008.

[42] D. V. Pozdyshev, A. A. Zharikova, M. V. Medvedeva, and V. I. Muronetz, “Differential Analysis of A-to-I mRNA Edited Sites in Parkinson’s Disease,” Genes, vol. 13, no. 1, p. 14, Dec. 2021, doi: 10.3390/genes13010014.

[43] V. Drouet and S. Lesage, “Synaptojanin 1 Mutation in Parkinson’s Disease Brings Further Insight into the Neuropathological Mechanisms,” BioMed Res. Int., vol. 2014, pp. 1– 9, 2014, doi: 10.1155/2014/289728.

[44] S. Tiwari, A. Singh, P. Gupta, and S. Singh, “UBA52 Is Crucial in HSP90 Ubiquitylation and Neurodegenerative Signaling during Early Phase of Parkinson’s Disease,” Cells, vol. 11, no. 23, p. 3770, Nov. 2022, doi: 10.3390/cells11233770.

[45] N. Khayer, M. Mirzaie, S.-A. Marashi, and M. Jalessi, “Rps27a might act as a controller of microglia activation in triggering neurodegenerative diseases,” PLOS ONE, vol. 15, no. 9, p. e0239219, Sep. 2020, doi: 10.1371/journal.pone.0239219.

[46] “parkinson’s disease related genes - GeneCards Search Results.” https://www.genecards.org/search/keyword?querystring=parkinson%27s%20%20disease&startPage=296&pageSize=25&sort=Symbol&sortDir=Ascending (accessed Sep. 19, 2023).

[47] N. Ball, W.-P. Teo, S. Chandra, and J. Chapman, “Parkinson’s Disease and the Environment,” Front. Neurol., vol. 10, p. 218, Mar. 2019, doi: 10.3389/fneur.2019.00218.

[48] S. Iannaccone et al., “In vivo microglia activation in very early dementia with Lewy bodies, comparison with Parkinson’s disease,” Parkinsonism Relat. Disord., vol. 19, no. 1, pp. 47–52, Jan. 2013, doi: 10.1016/j.parkreldis.2012.07.002.

[49] P. Bossù, G. Spalletta, C. Caltagirone, and A. Ciaramella, “Myeloid Dendritic Cells are Potential Players in Human Neurodegenerative Diseases,” Front. Immunol., vol. 6, Dec. 2015, doi: 10.3389/fimmu.2015.00632.

[50] X. Su and H. J. Federoff, “Immune Responses in Parkinson’s Disease: Interplay between Central and Peripheral Immune Systems,” BioMed Res. Int., vol. 2014, pp. 1–9, 2014, doi: 10.1155/2014/275178.

[51] E. C. Hirsch and D. G. Standaert, “Ten Unsolved Questions About Neuroinflammation in Parkinson’s Disease,” Mov. Disord., vol. 36, no. 1, pp. 16–24, Jan. 2021, doi: 10.1002/mds.28075.

[52] L. Muñoz-Delgado et al., “Peripheral inflammatory immune response differs among sporadic and familial Parkinson’s disease,” Npj Park. Dis., vol. 9, no. 1, p. 12, Jan. 2023, doi: 10.1038/s41531-023-00457-5.

[53] M. Alves, D. Caldeira, and J. J. Ferreira, “Blood pressure variability in Parkinson’s Disease patients – Case control study,” Clin. Park. Relat. Disord., vol. 8, p. 100191, 2023, doi: 10.1016/j.prdoa.2023.100191.

[54] B. S. Desai, A. J. Monahan, P. M. Carvey, and B. Hendey, “Blood–Brain Barrier Pathology in Alzheimer’s and Parkinson’s Disease: Implications for Drug Therapy,” Cell Transplant., vol. 16, no. 3, pp. 285–299, Mar. 2007, doi: 10.3727/000000007783464731.

[55] M. T. Fuzzati-Armentero, S. Cerri, and F. Blandini, “Peripheral-Central Neuroimmune Crosstalk in Parkinson’s Disease: What Do Patients and Animal Models Tell Us?,” Front. Neurol., vol. 10, p. 232, Mar. 2019, doi: 10.3389/fneur.2019.00232.

[56] “HLA-F major histocompatibility complex, class I, F [Homo sapiens (human)] - Gene - NCBI.” https://www.ncbi.nlm.nih.gov/gene/3134 (accessed Sep. 24, 2023).

[57] “IRF1 interferon regulatory factor 1 [Homo sapiens (human)] - Gene - NCBI.” https://www.ncbi.nlm.nih.gov/gene/3659 (accessed Sep. 24, 2023).

[58] J. Emile, J. Truelle, A. Pouplard, and D. Hurez, “[Association of Parkinson’s disease with HLA-B17 and B18 antigens],” Nouv. Presse Médicale, vol. 6, p. 4144, Jan. 1978.

[59] E. Yu et al., “Fine mapping of the HLA locus in Parkinson’s disease in Europeans,” NPJ Park. Dis., vol. 7, p. 84, Sep. 2021, doi: 10.1038/s41531-021-00231-5.

[60] T. Taniguchi, K. Ogasawara, A. Takaoka, and N. Tanaka, “IRF family of transcription factors as regulators of host defense,” Annu. Rev. Immunol., vol. 19, pp. 623–655, 2001, doi: 10.1146/annurev.immunol.19.1.623.

[61] F. Mu, X. Chen, X. Du, Q. Jiao, M. Bi, and H. Jiang, “[Regulatory mechanism of interferon regulatory factor 1 by α-synuclein in mouse Parkinson’s disease model],” Nan Fang Yi Ke Da Xue Xue Bao, vol. 41, no. 11, pp. 1641–1648, Nov. 2021, doi: 10.12122/j.issn.1673-4254.2021.11.07.

[62] “RPS28 ribosomal protein S28 [Homo sapiens (human)] - Gene - NCBI.” https://www.ncbi.nlm.nih.gov/gene/6234 (accessed Sep. 24, 2023).

[63] U. Stelzl, S. Connell, K. H. Nierhaus, and B. Wittmann_Liebold, “Ribosomal Proteins: Role in Ribosomal Functions,” in eLS, John Wiley & Sons, Ltd, Ed., 1st ed.Wiley, 2001. doi: 10.1038/npg.els.0000687.

[64] D. N. Wilson et al., “Protein synthesis at atomic resolution: mechanistics of translation in the light of highly resolved structures for the ribosome,” Curr. Protein Pept. Sci., vol. 3, no. 1, pp. 1–53, Feb. 2002, doi: 10.2174/1389203023380846.

[65] P. Garcia-Esparcia et al., “Altered machinery of protein synthesis is region- and stage-dependent and is associated with α-synuclein oligomers in Parkinson’s disease,” Acta Neuropathol. Commun., vol. 3, no. 1, p. 76, Dec. 2015, doi: 10.1186/s40478-015-0257-4.

[66] Y. Jang et al., “Mass Spectrometry–Based Proteomics Analysis of Human Substantia Nigra From Parkinson’s Disease Patients Identifies Multiple Pathways Potentially Involved in the Disease,” Mol. Cell. Proteomics, vol. 22, no. 1, p. 100452, Jan. 2023, doi: 10.1016/j.mcpro.2022.100452.

[67] D. Chicco and G. Jurman, “The advantages of the Matthews correlation coefficient (MCC) over F1 score and accuracy in binary classification evaluation,” BMC Genomics, vol. 21, no. 1, p. 6, Dec. 2020, doi: 10.1186/s12864-019-6413-7.

[68] D. Chicco and G. Jurman, “The Matthews correlation coefficient (MCC) should replace the ROC AUC as the standard metric for assessing binary classification,” BioData Min., vol. 16, no. 1, p. 4, Feb. 2023, doi: 10.1186/s13040-023-00322-4.

[69] “GitHub: Let’s build from here,” GitHub. https://github.com/ (accessed Sep. 19, 2023).

[70] A. Bhattacharjee, “PitArray (ParkInson’s disease detection Tool using blood-based microArray data).” Sep. 12, 2023. Accessed: Sep. 19, 2023. [Online]. Available: https://github.com/Arittra95/PitArray

[71] E. Bisong, “Google Colaboratory,” in *Building Machine Learning and Deep Learning Models on Google Cloud Platform: A Comprehensive Guide for Beginners*, E. Bisong, Ed., Berkeley, CA: Apress, 2019, pp. 59–64. doi: 10.1007/978-1-4842-4470-8_7.

