## Supplementary figures and images for "A Non-invasive Detection of Parkinson’s Disease using PitArray: An Integrative Meta-Analysis and Machine Learning Approach"

### Supplementary Figure 1

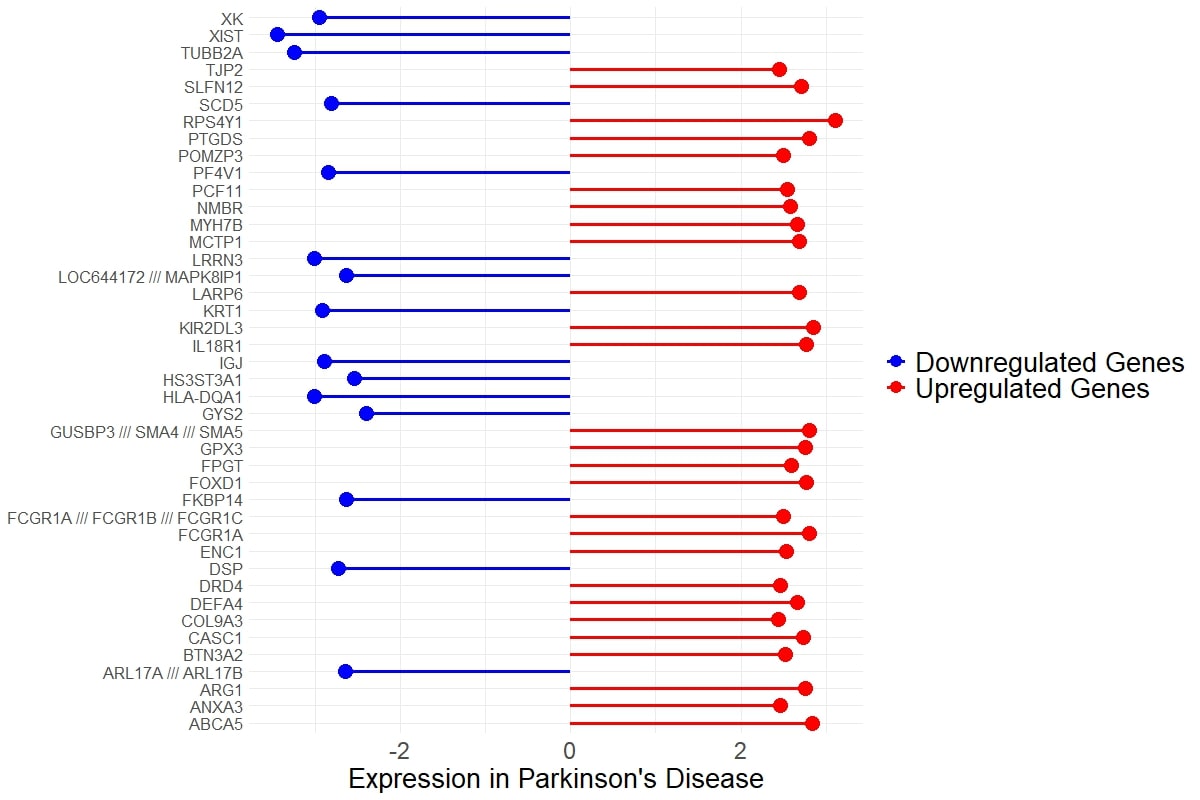
